# Dissecting DNA-mismatch-repair-driven mutational processes in human cells

**DOI:** 10.1101/2025.02.10.637460

**Authors:** Michel Owusu, Jörg Menche, Joanna Loizou, Donate Weghorn

## Abstract

Mismatch repair (MMR) is an essential DNA repair pathway that preserves genome stability by correcting replication errors and other DNA lesions. MMR deficiency (MMRd) leads to increased mutation rates and microsatellite instability, and contributes to tumorigenesis in multiple cancer types, yet the full spectrum of mutational patterns caused by MMRd remains incompletely defined. Here, using CRISPR–Cas9 knockouts in human isogenic cell lines, we systematically profile mutations arising in MMR-deficient cells. We observe a marked increase in mutation burden and the appearance of established MMRd-associated mutational signatures. Unexpectedly, we also detect signatures SBS57 and DBS14 and connect them to germline single-nucleotide polymorphisms that co-occur with MMR-driven indels. In addition, we provide experimental genomic evidence supporting a role of MMR in preventing mutations underlying the widespread SBS1 signature, and we link MMR deficiency to the indel signature ID12, which currently lacks a known aetiology. Finally, comparing *in vitro* MMRd profiles from rapidly dividing cells without exogenous mutagen exposure to *in vivo* tumour genomes that reflect lifelong mutation accumulation reveals pronounced differences in signature composition, underscoring the context dependence of MMR-linked mutagenesis. Together, our results broaden the known mutational footprint of MMR deficiency and illuminate previously uncharacterised mechanisms relevant to cancer development.

## Introduction

Mutations are an unavoidable consequence of unrepaired damage to DNA. A particularly important DNA repair pathway is mismatch repair (MMR). MMR was originally discovered in the context of base-base mismatches [1], i.e., incorrect nucleotide pairings introduced by DNA polymerases during replication, but we now know that MMR functions in many other contexts of DNA damage repair. These include homologous recombination and the recognition of alkylated DNA, oxidised DNA and interstrand crosslinks [2]. Recently, MutS*α*-driven MMR activity during CpG deamination repair was demonstrated using sequencing data from healthy and tumour tissues [3, 4], expanding the catalogue of MMR activities [5]. Deficiency of the MMR machinery can lead to an increase in the rate of single-base substitutions (SBSs) and small insertions and deletions (indels), the latter caused by polymerase slippage, resulting in the cellular phenotype of microsatellite instability (MSI). Several disease phenotypes are associated with MMR deficiency, including hereditary nonpolyposis colorectal carcinoma (HNPCC), also known as Lynch syndrome, which is mediated by monoallelic mutations in MMR genes [6], constitutional mismatch repair deficiency (CMMRD), which is mediated by biallelic MMR mutations [7, 8, 9], and other MSI tumours mediated by somatically acquired MMR deficiency [10]. Extraction of mutational signatures has identified a variety of independent mutational processes associated with defective mismatch repair [11]. The fact that there is more than one such signature suggests that MMR is a composite process driven by many different interacting players, but their interplay is currently unknown.

Here, we generated CRISPR-Cas9-mediated knockouts of a selection of MMR genes in isogenic human HAP1 cells, followed by whole-genome sequencing, to investigate the mutational profiles in cells without functional MMR. In addition to an expected increase in overall mutational burden, we identified several mutational signatures associated with MMR deficiency (MMRd). Analysis of the dependence of mutational signatures on replication time revealed an additional unexpected process enriched in early-replicating regions. We show that this process, SBS57, is driven by the synergistic effects of pre-existing germline single-nucleotide polymorphisms (SNPs) and MMR-related indels on sequencing read alignment in MMR-deficient cells, and link it to the tensor signature TS27 [12]. In a direct comparison of *in vitro* and *in vivo* MMR deficiency, we further uncover differences in MMRd-associated signatures between cell growth at high division rates without exogenous mutagens (cell culture experiments) and malignant transformation over a patient’s lifetime (cancer tumours). Finally, we provide experimental genomic evidence supporting the notion that MMR contributes to preventing SBS1 mutations arising from 5-methylcytosine deamination, a relationship previously inferred from sequencing data from human tissues. Overall, our results shed light on new aspects of MMR deficiency, including the driving mechanisms underlying signatures of previously unknown aetiology.

## Results

### CRISPR-Cas9-mediated knockout of canonical mismatch repair genes causes a variety of mutational signatures

To investigate the role of DNA mismatch repair, we generated isogenic human HAP1 cells with individual gene knockouts (KOs) using CRISPR-Cas9-mediated frameshift deletions. We selected canonical MMR genes, those directly involved in MMR (*MLH1*, *MSH2* and *MSH6*), as well as genes indirectly associated with the pathway (**Figure 1A**). These include both established interactors (*ARID1A* [13], *EXO1* [14], *SETD2* [15]) and potential interactors of MMR. Those are *NSD1*, a regulator of H3K36me3, a histone mark involved in the recruitment of MutS*α* [16], and *SMARCA4*, a core component of the SWI/SNF chromatin remodeling complex, which plays a central role in various cellular processes, including DNA repair. After a growth period of three months in culture, single cells were clonally expanded in three replicates (**Figure 1B**). Comparison with the clonal expansions of the starting cells allowed us to determine only mutations that had arisen *de novo*.

**Figure 1:**
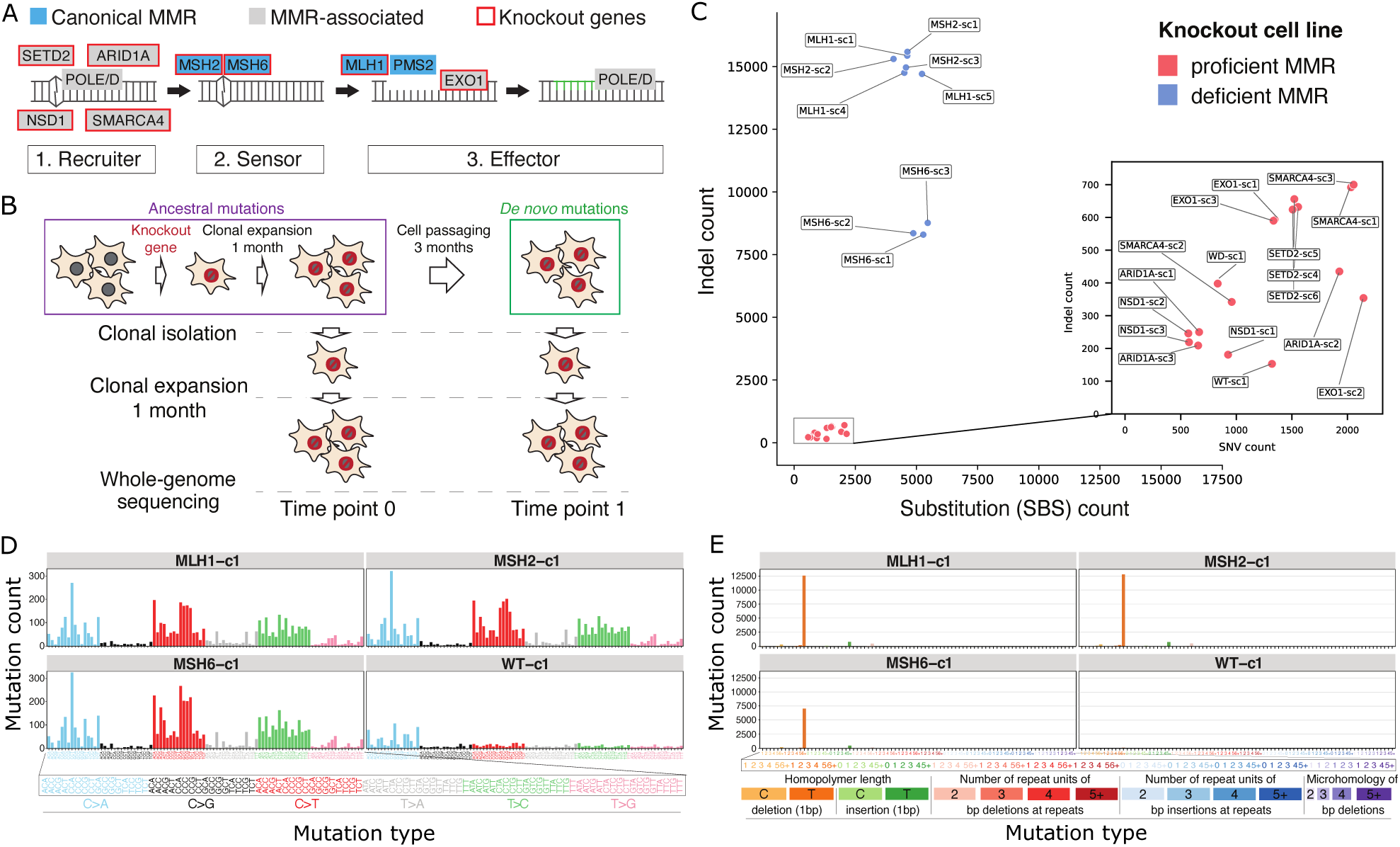
Experimental strategy and observed mutational profiles. (A) Genes selected for *in vitro* CRISPRCas9-mediated knockout (KO). (B) Experimental workflow: Single KO cells were clonally expanded for one month. Cells reflecting the initial genetic makeup (*t*_0_) were obtained after clonal expansion starting from single cells after the first month. The remaining cells were cultured for another three months, followed by clonal expansion (*t*_1_). Mutations present in *t*_1_ but not *t*_0_ were considered *de novo*. (C) Counts of *de novo* singlebase substitutions and small insertions and deletions (indels) are shown for all KOs (including technical replicates) and wild-type cells (WT) after data curation. (D,E) Mutational profiles of *MLH1*, *MSH2*, *MSH6* KO and WT cells showing single-base substitutions (D) and indels (E).

We found that three gene knockouts caused a repeated increase in the observed number of singlebase substitutions and indels compared to wild type (WT): *MLH1*, *MSH2* and *MSH6* (“MMRd KOs”; **Figure 1C, Supplementary Figure 1**). Conversely, the *SETD2* and *EXO1* knockouts had a more limited effect on the number of mutations, consistent with previous reports [2]. Notably, none of the other knockouts showed a consistent increase in mutation count in our experimental setup over the course of three months, suggesting that these genes have functional backups or are mutagenic when exposed to exogenous stressors.

By classifying mutations according to their sequence context and the alternate base or indel type, we categorised the observed mutations into mutational profiles (**Figure 1D-E, Supplementary Figures 2-3**). *MLH1*, *MSH2* and *MSH6* KOs exhibited similar mutational profiles consistent with previous reports [17, 18, 19]. Next, we refitted the SBS and indel (ID) mutational profiles of all KO cell lines to known SBS and ID mutational signatures from the COSMIC signature catalogue [11] (**Figure 2A**). As expected, each sample in our experiment had a sizeable (*>*5%) contribution of signature SBS18, a signature frequently observed under experimental cell culture conditions and associated with oxidative DNA damage [17, 18]. Several substitution signatures were found exclusively in the cells with the MMRd mutational profile: SBS6, SBS14, SBS15, SBS20, SBS21, SBS26 and SBS44, which are all the seven COSMIC signatures currently associated with defective MMR [11]. SBS44 also has the highest individual contribution of all signatures in the MMRd cells.

**Figure 2:**
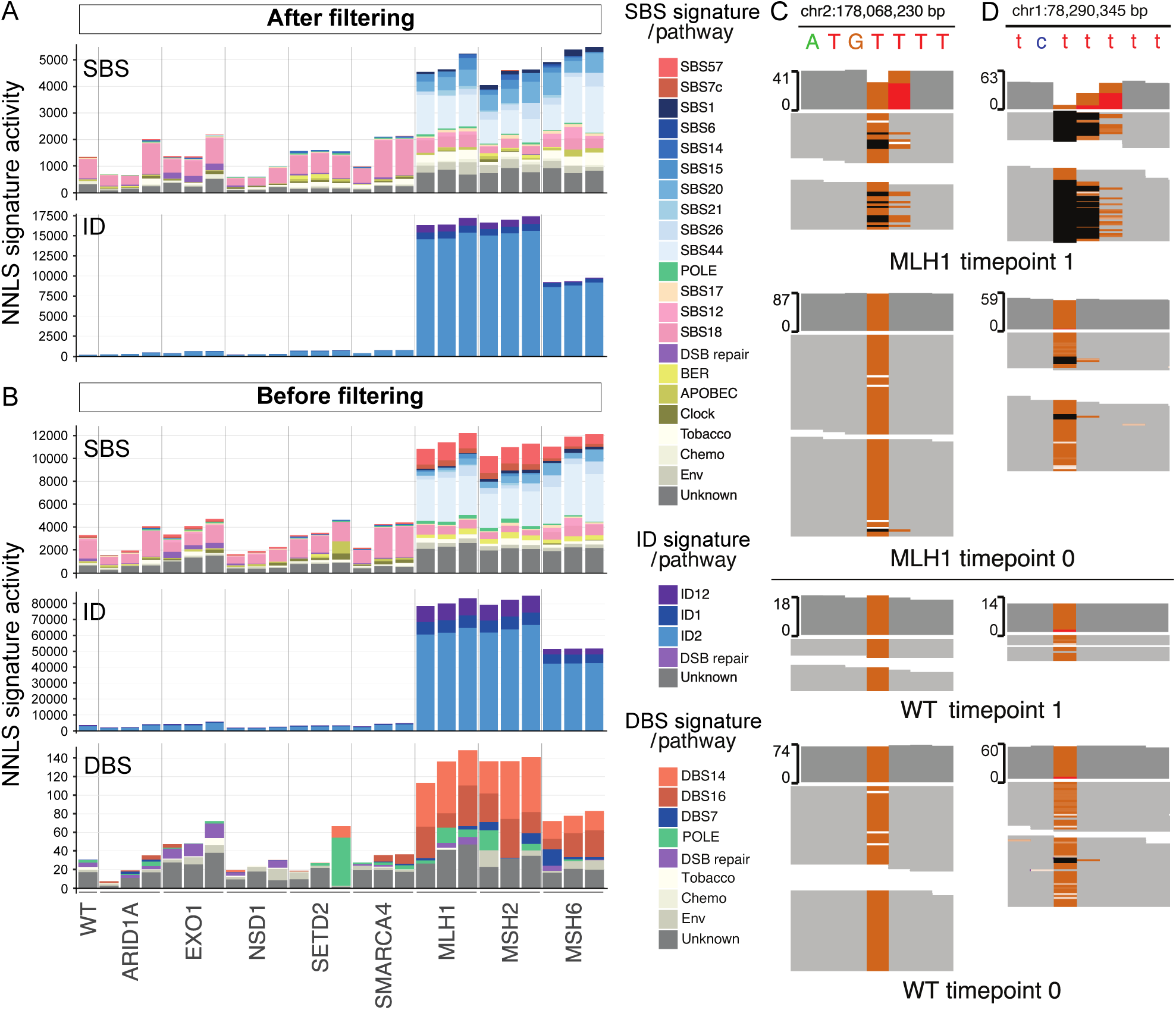
Mutational signature refitting identifies MMR-associated signatures and SBS57 in knockout cells. Mutational profiles of the experimental cell lines were refitted to COSMIC mutational signatures v3.4 [11]. Known MMR-associated signatures and SBS1 are shown in shades of blue. Other signatures are categorised by biological pathways (**Methods**). (A) SBS and ID signatures for all KO replicates and WT cells for consensus calls after applying the repeat-masker filter. Note that after filtering less than 100 DBSs remained across all samples, so signature decompositions are not shown. (B) SBS, ID and DBS signatures for all KO replicates and WT cells without applying additional mutation filters (i.e., Strelka2 calls in mappable regions). All signatures obtained from non-negative least-squares refitting to COSMIC v3.4 signatures. (C-D) Representative snapshots of aligned reads around a typical SBS57-associated mutation (TTT*>*TGT; C) and a typical DBS14-associated mutation (TT*>*GG; D) from the Integrative Genomics Viewer (IGV). Grey indicates the reference genome. The arrows indicate germline T*>*G SNPs present in all cellular replicates (*MLH1* and WT) and time points (*t*_0_ and *t*_1_). Black indicates a deletion. Note that SNPs do not appear as heterozygous because cells come from an almost fully haploid cell line (HAP1). SBS=single-base substitution, ID=indel, DBS=double-base substitution, POLE=polymerase *ε*, DSB=double-strand break, BER=base-excision repair, Clock=clock-like, Chemo=chemotherapy, Env=environment.

At first sight, the sizeable amounts of SBS14 and SBS20 should indicate alterations of polymerase epsilon (POLE) and delta (POLD) function, respectively, concomitant with MMRd. However, we neither found genetic POLE or POLD variants in our cells, nor do we observe the excess of SBS10a/b that would be expected for POLE deficiency. SBS14 is characterised by mutational peaks at XCT*>*XAT (X*2*{A,C,G,T}), while SBS20 exhibits a strong CCT*>*CAT peak, followed by a slightly less pronounced excess of CCC*>*CAC mutations. These two features of SBS20 were also observed in a recent experiment using human iPSCs and it was hypothesised that they signalled MMR involvement in repairing 8-oxo-G:A mismatches, rather than true presence of POLD deficiency [19]. This is compatible with previous indications that MMR excises incorporated 8-oxodGMP from newly synthesized daughter DNA [20, 21].

While MMRp cells had no detectable SBS1 signature, we found substantial amounts of this signature in the MMR KOs, mainly in the *MSH2* and *MSH6* KOs. This is consistent with observations from DNA sequencing data of cancer and healthy tissue suggesting a link between the repair of deaminated 5-methylcytosine and MutS*α* [5, 3, 4]. Thus, our isogenic knockout experiments provide direct experimental support for this link.

As for indels, among the largest contributions in the MMRd samples came from ID1 and ID2, the two signatures associated with DNA polymerase slippage during replication causing 1-bp insertions or deletions in thymine homopolymers, consistent with the lack of repair of these variants by MMR. The predominance of ID2 is characteristic of MMRd [11]. Suprisingly, we also identified a large number of ID12 mutations in MMR KOs. This indel signature is dominated by 2-bp deletions in repeat-rich regions and has no known aetiology. Our results suggest that ID12 is associated with MMRd, consistent with recent reports of MMRd tumours from mice [22].

Although *EXO1* and *SETD2* loss caused only modest increases in total mutation burden, both KOs showed recurrent non-MMR mutational patterns. Across all three *EXO1* clones, the SBS group with putative links to double-strand break repair (comprising SBS3, SBS8 and SBS39) showed a 7.0-fold enrichment (two-sided exact permutation P=0.006). EXO1 contributes to extended DNA end resection and promotes RPA and RAD51 recruitment during homologous recombination [23, 24]. While SBS3 is a well-known homologous-recombination-deficiency-(HRD)-associated signature, SBS8 has been linked to uncorrected errors in late-replicating genomic regions and to co-occurrence with SBS3 and SBS39 [25, 26]. The relaxed-filter double-base-substitution profiles provided a concordant signal: COSMIC DBS13, an HRD-associated signature [27], accounted for a 6.9-fold enrichment in *EXO1* KOs (P=0.006). At the same time, HRD-associated indel signatures ID8 and ID6 were not statistically significantly enriched. Together, these findings are consistent with altered double-strandbreak-repair pathway usage following *EXO1* loss, but do not indicate a complete *BRCA*-like HRD phenotype. *SETD2* knockout was most clearly associated with SBS36, showing a 24-fold enrichment (P=0.012). SBS36 has been experimentally linked to defective processing of 8-oxoG-associated lesions by the base excision repair (BER) pathway [19]. Consistent with this interpretation, SETD2-null cells have been shown to accumulate 8-oxoG adducts and to exhibit impaired oxidative-damage signalling [28]. Thus, our findings extend the established biochemical role of SETD2 in 8-oxoG processing to a genome-wide mutational phenotype.

### Single-base-substitution signature SBS57 is elevated in MMR knockouts

The results shown in **Figure 1** and **2A** were obtained after applying a strict filter to the mutation calls (**Methods**). Specifically, in addition to generating consensus calls from the output of two mutation callers (Strelka2 and MuTect2), we filtered out variants in regions with repeats after using a baseline mappability filter. We applied the additional consensus filter and repeat masker after detecting a large contribution of the unexpected signature SBS57 with weights of 5-15% in MMRd samples but not in MMRp samples (**Figure 2B**). SBS57 is classified as a “possible sequencing artifact” in the COSMIC database. Based on mutations called with one caller (Strelka2), SBS57 was recently hypothesised to be associated with chemical DNA damage due to formalin fixation (FFPE), although the same study also found the signature in fresh-frozen tumour samples [29]. Another study reported SBS57 in different cell types after irradiation [30]. We found that SBS57 in MMRd cells shows a clear dependence on replication time, with high average mutation numbers in the earliest replication quantile that gradually decrease towards the last replication quantile (**Supplementary Figure 4**).

Since neither sequencing technologies nor FFPE are expected to introduce such a strong correlation between mutational signatures and replication time, we hypothesised that the mechanism underlying SBS57 had a different basis. SBS57 is characterised by spikes of TTT*>*TCT and TTT*>*TGT mutations. When we examined these “SBS57-like” mutations in more detail using the Integrative Genomics Viewer (IGV), we found that the read distribution around the SBS57-like mutations has very specific features: (1) a germline C or G SNP within a few base pairs of the site of the mutated T, (2) a small *de novo* deletion within a few base pairs of the mutated site, and (3) the mutated site is located at or near the boundary of a thymine repeat region. We discovered that the combination of these three features resulted in erroneous shifts of the germline SNPs in the alignment of reads containing both the SNP and the deletion, leading to false-positive SBS57-like mutations (**Figure 2C**). In addition to SBS57, the same deletion- and SNP-associated alignment error produced DBS14-like TT*>*NN calls in the unfiltered MMRd cell data (**Figure 2D**). Although DBS14 has previously been linked to read misalignment and shown to co-occur with SBS57 in MMRd tumours [31], our results identify a specific local alignment mechanism capable of generating both signatures.

### SBS57 is detected in MMR-deficient and other cancer tumours

SBS57 was originally described in cancer cells without an established link to MMR. We therefore investigated the relationship between SBS57 and MMR in metastatic tumours from the Hartwig Medical Foundation (HMF; mutation calling described in ref. [32]), restricting the analysis to samples with at least 1,000 substitutions. Using a probabilistic criterion for reliable detection of SBS57 (**Methods, Supplementary Figure 5**), we identified 223 tumour samples with SBS57. We performed principal component analysis of the COSMIC v3.4 SBS signature weights, excluding SBS57 and signatures with no variation across the selected tumours, followed by k-means clustering based on the first two principal components. This separated the tumours into three groups: 42 MMRd-like, 87 SBS17-like and 94 Other tumours (**Figure 3A**). Samples with silhouette scores ≤0.631 were considered less confidently assigned and were excluded from the downstream cluster-specific analyses, leaving 29 MMRd-like, 47 SBS17-like and 66 Other tumour samples. The MMRd-like group was enriched for canonical MMRd-associated signatures, whereas the SBS17-like group was dominated by SBS17a and SBS17b (**Figure 3B**). Intriguingly, an association between SBS17 and MMR deficiency was found previously [3].

**Figure 3:**
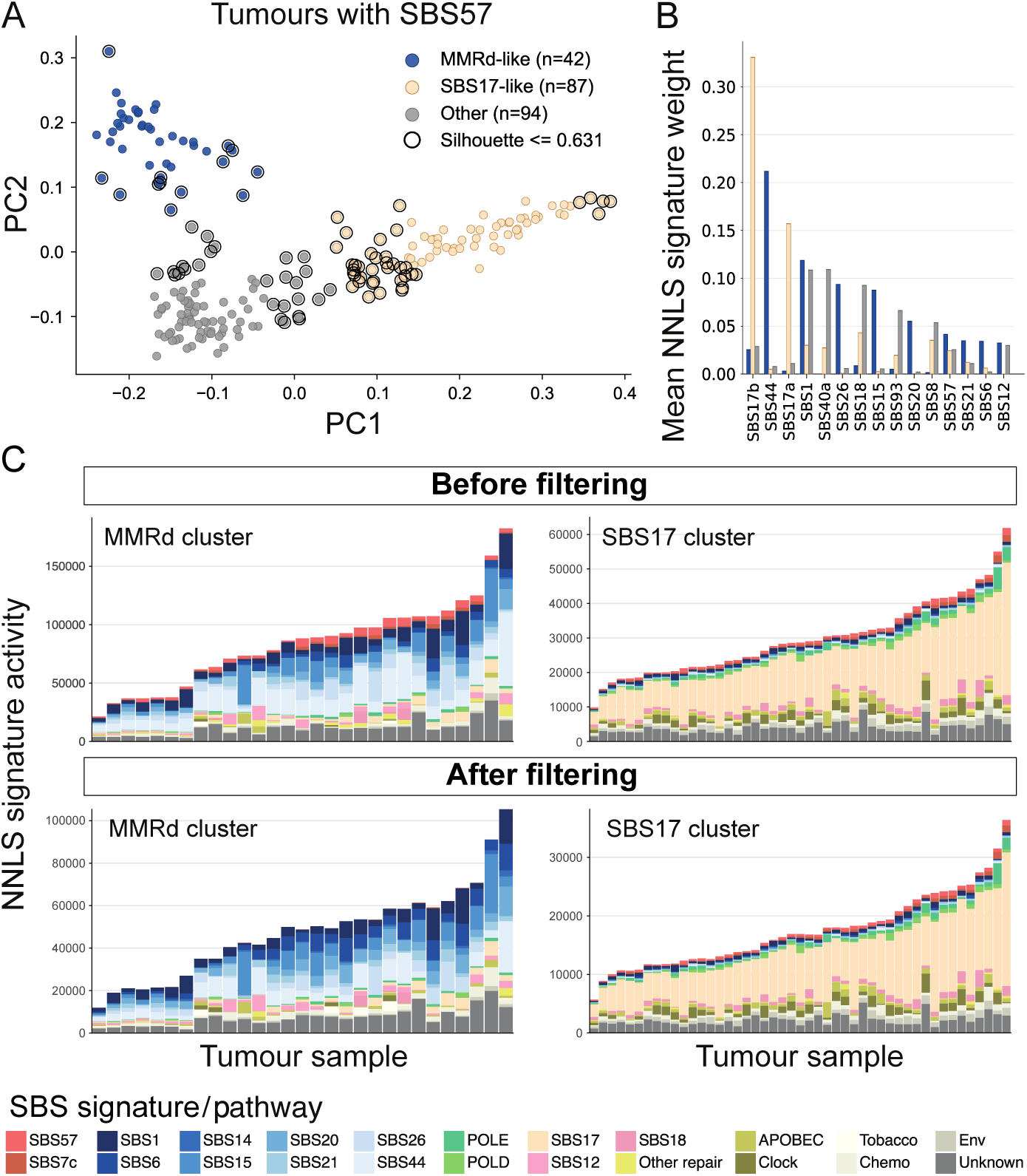
Identification of SBS57-positive tumours. (A) First two principal components (PCs) of the COSMIC signature weights of 223 SBS57-positive tumour samples from the HMF cohort (4709 samples), coloured by k-means cluster assignment. (B) Most common signatures across tumour cluster identities. (C) Single-base-substitution (SBS) signature refitting results from SBS57-containing tumour samples from the MMRd-like and the SBS17-like cluster (see **Supplementary Figure 5** for Other cluster) obtained from non-negative leastsquares refitting to COSMIC v3.4 signatures. Top: Mutation calling without extra filters (i.e., HMF-provided Strelka calls restricted to mappable genomic regions). Bottom: Mutation calling with applying the additional filtering scheme (i.e., exclusion of variants in repeat-masked regions and/or within *±*6 bp of small deletions). HMF=Hartwig Medical Foundation.

As in the KO cells, we next assessed how variant filtering affected SBS57 detection in tumours. We removed variants in non-mappable and repeat-masked sites and additionally applied a deletionbased variant filter, as described in the **Methods**. In the MMRd-like tumours, the combination of these filters completely eliminated SBS57, recapitulating the effect observed in the MMRd KO cells (**Figure 3C**). Applying the repeat-masker and mappability filters alone, without the additional deletion-based filtering step, left a small residual SBS57 contribution (**Supplementary Figure 6**), suggesting that the consensus mutation calling filtered this residual out in the MMRd cell line analysis. By contrast, SBS57 persisted after the same combined filtering in the SBS17-like and Other tumour groups, arguing against SBS57 arising through the repeat- and deletion-associated process identified in MMRd cells in tumours without an MMRd phenotype.

Besides the COSMIC mutational signatures, another powerful mutational signature extraction framework was proposed by Vöhringer *et al.* (2021) [12]. Refitting the KO samples to this alternative set of mutational signatures, we found that one signature recapitulated the SBS57 results: TS27 (**Supplementary Figure 7A**). TS27 has a strong replication strand bias, but otherwise an unknown aetiology [12]. The difference between SBS57 and TS27 suggests that SBS57 lacks components in the T*>*A category, especially ATT*>*A and TTT*>*A. We attribute this to the unexpected identification of SBS7c, one of the four normally co-occurring signatures associated with UV damage, in almost all of the KO samples and the MMRd tumours, which also disappeared after applying the filtering scheme (**Figures 2B** and **3C**). Therefore, we propose that SBS57 identification in the MMRd context is a proxy for a mutational process more accurately represented by TS27 (which has no direct analogue in the COSMIC database). This is also supported by the almost perfect co-occurrence (Jaccard index = 0.83) of TS27 with two MMRd-associated tensor signatures in the HMF cohort (TS16, similar to SBS6 and SBS14, and TS26, resembling SBS15) [12]. TS16 is also the most common tensor signature identified in the MMRd cells after TS27. Finally, when we applied the filtering scheme TS27 completely disappeared, as before SBS57 (**Supplementary Figure 7B**). Of note, TS27 also shows the typical spikes in the CpG*>*TpG mutation categories otherwise associated with SBS1, which we identified in the MMRd samples but not in the MMRp cells.

### MMR deficiency footprints *in vitro* and *in vivo*

While the knockout cell culture experiments serve as a system to study the effects of MMR deficiency under controlled conditions, they are fundamentally different from the dynamics in a growing cancer tumour. First, experimental conditions cause increased exposure to oxidative damage (SBS18) in cell culture, but other tumorigenic mutagens are lacking. Second, cell division rates in culture are greatly increased due to the abundance of nutrients and space (serial dilutions). At the same time, the total growth time is only three months, compared to months or years experienced by a growing tumour. We hypothesised that the differences in growth rate and total growth time would lead to differences between MMR-associated signatures. Specifically, since mutations are the product of the combined effect of DNA damage and (imperfect) repair, we expect that mutational processes that depend on damage accumulation (on an absolute time scale or between cell divisions) would be more pronounced in tumours, whereas strictly replication-driven processes should have a higher relative impact in cell culture.

The controlled MMRd KO system and MMRd tumours exhibited clearly distinct mutational profiles (**Figure 4A**). To determine whether these differences reflected changes in the underlying mutational processes, we compared their mutational signature compositions (**Figure 4B**). The most striking signature difference was the almost complete absence of SBS6 in the MMRd KO cells, suggesting that this signature is caused by the accumulation of damage over extended periods of time. Several other processes were also relatively enriched in the tumours: SBS1, SBS15, SBS21, and SBS26. Conversely, the relative proportion of SBS44 was 3.7-fold greater in the MMRd KO cells (0.25 vs. 0.07, which holds also after removing SBS6 and the non-MMRd-associated signatures), suggesting that this signature could be affected by the differences in microenvironment between tumours and cell cultures or caused by a replicative process. The latter is consistent with the replication strand bias observed for the major SBS44-associated mutation types in the strand-stratified mutational profiles (**Figure 4A**).

**Figure 4:**
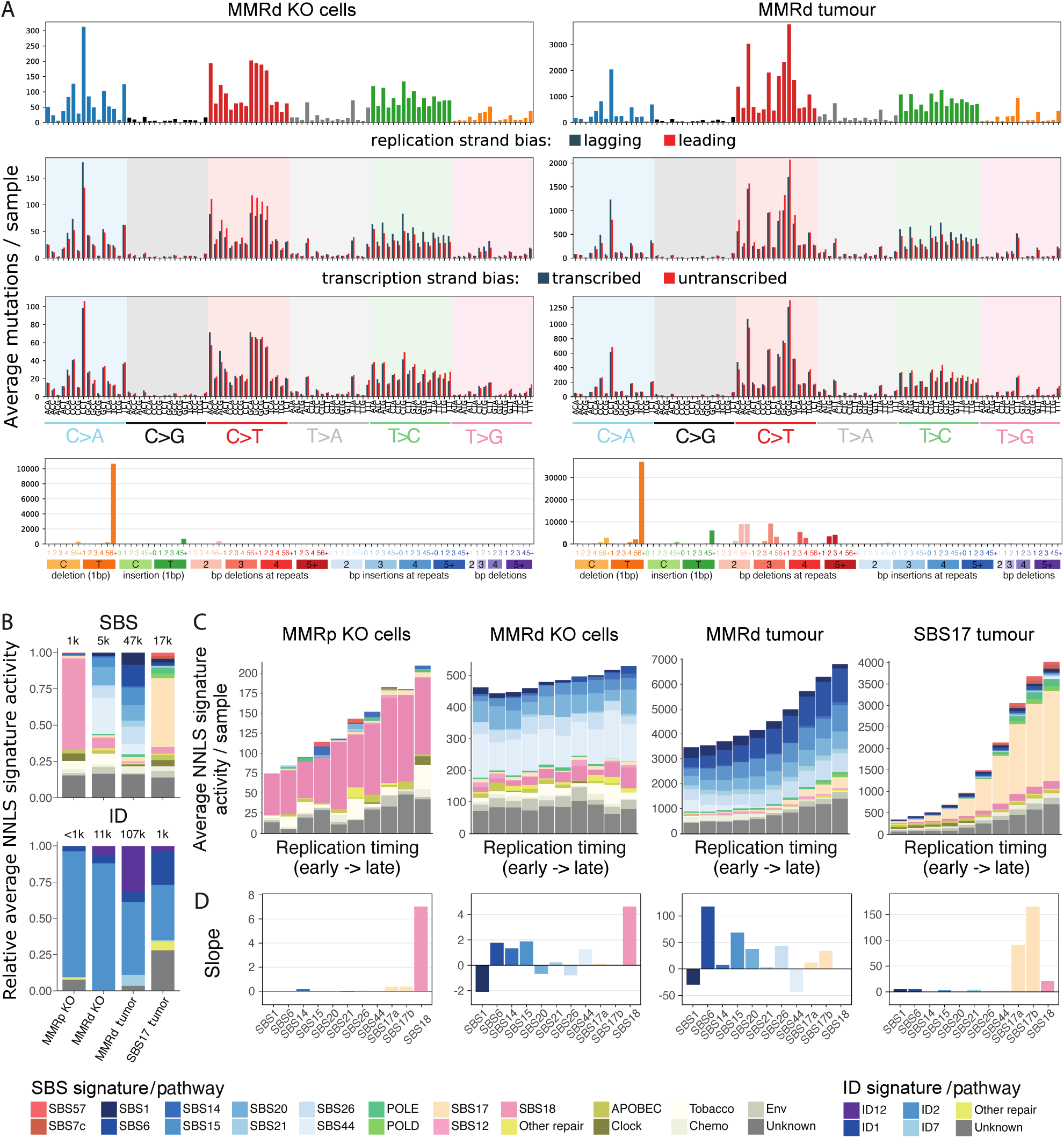
Differences between *in vitro* and *in vivo* MMR deficiency. (A) Comparison of MMRd KO cell cultures (left) and MMRd-like cancer tumours (right). Top to bottom: Average MMRd SBS mutational profiles, SBS replication strand bias, SBS transcription strand bias and MMRd indel mutational profiles. (B) Average COSMIC mutational signature weights in MMRp cells (defined in **Methods**), MMRd cells (MLH1, MSH2 and MSH6) and the MMRd-like and SBS17-like cancer tumour clusters for SBSs (top) and indels (bottom). (C) Sequence-context-opportunity-corrected replication time dependence of signature-associated mutation counts in MMRp cells, MMRd cells, MMRd-like tumours and SBS17-like tumours, obtained from non-negative least-squares refitting to COSMIC v3.4 signatures. (D) Slopes of linear model fits to the data shown in (C). All data are shown after applying the strict filtering scheme. SBS=single-base substitution.

We found SBS1 both *in vitro* and *in vivo*, with a higher relative weight in tumours. Although SBS1 is referred to as a clock-like signature, i.e., a signature that increases with time (age), it was found to be age-independent in post-mitotic cells, linking it to cell division, compatible with a role of MMR in repair of deaminated 5-methylcytosine [33, 5]. At the same time, the damage underlying SBS1 itself is not directly associated with replication and canonically repaired by base excision. Together, this implies that SBS1 is driven both by high replication rates that convert unrepaired deaminated CpGs into TpGs and by intermittent damage accumulation that increases over time.

The tumours showed a decrease of about 43% in ID2 proportion, evidenced by a ratio of *≥*6-homopolymer-associated 1-bp T deletions to SNVs of around 2.2 in the filtered mutations from MMRd cells (5.5 when using only Strelka2 calls and including repeat-masked regions, **Supplementary Figure 1**) that decreased to only 0.8 in MMRd tumours with repeat-masked Strelka mutation calls [32] (**Figure 4A**). Notably, ID7, a known MMRd-associated indel signature, was only present in MMRd tumours, suggesting an association with damage accumulation, rather than replication, or exogenous mutagens not present in our cell cultures. As before for the MMRd cells, also the MMRd tumours showed ID12, independently supporting an association of ID12 with defective mismatch repair.

Finally, we examined how mutagenesis in the two MMRd systems varied with replication timing (RT). Consistent with pan-cancer analyses of mutational topography [34], the opportunity-corrected, artefact-filtered data showed enrichment of mutations in late-replicating regions in both MMRd cells and tumours (**Figure 4C**). A clear excess in early-replicating regions was observed only before filtering and was attributable to SBS57-associated artefacts, indicating that it does not represent a biological MMRd mutational process (**Supplementary Figure 4**).

Individual MMR-associated signatures exhibited distinct RT profiles (**Figure 4D, Supplementary Figures 8-12**). In both cells and tumours, SBS1 was enriched in early-replicating regions, whereas SBS6, SBS14 and SBS15 increased towards later-replicating regions. SBS44 showed contrasting behaviour between the two systems. In the cell lines, it was detected across all RT bins and increased slightly towards late replication. In tumours, by contrast, SBS44 was concentrated in early-replicating regions and was absent from the late-replicating bins. SBS21 showed little dependence on replication timing in either system. Together, these findings indicate that replication timing shapes MMRd mutagenesis in a signature- and context-dependent manner, with SBS44 in particular revealing marked differences between experimental cell systems and human tumours.

## Discussion

Using isogenic human HAP1 cells, we characterised the mutational consequences of mismatch-repair deficiency in a controlled genetic background. Previous CRISPR-based studies in human organoids, HAP1 cells and induced pluripotent stem cells established characteristic mutational phenotypes following disruption of MMR genes [17, 18, 19, 3]. Canonical MMR knockouts reproduced the expected MMRd-associated substitution and indel signatures, while also providing experimental support for MMR involvement in mutational processes not usually considered canonical MMR phenotypes. In particular, SBS1 accumulated after loss of MSH2 or MSH6, consistent with previous evidence that MutS*α* participates in repair of G:T mismatches caused by 5-methylcytosine deamination [4]. The recurrent presence of ID12 further suggests that MMR suppresses a broader class of repeat-associated deletions in addition to the well-established ID1/ID2 polymerase-slippage processes, consistent with observations in Msh2-deficient mice and constitutional MMR-deficiency tumours [22, 35].

We also identified a technical mechanism generating apparent SBS57 and DBS14 activity in MMRd cells. Nearby *de novo* deletions at thymine-repeat boundaries displaced pre-existing germline SNPs during read alignment, producing false-positive substitutions and double-base substitutions. This provides a specific sequence-level explanation for the previously reported association of DBS14 with read misalignment [31]. The persistence of SBS57 after filtering in SBS17-like and Other tumours, however, indicates that SBS57 is not uniquely caused by this artefact.

Comparison with MMRd-like tumours showed that MMR deficiency does not generate a fixed signature mixture. SBS44 was relatively prominent in the cell system, whereas SBS6, SBS15, SBS21 and SBS26 were more prominent in tumours, and ID7 was detected only in tumours. Replicationtiming profiles also differed between the two systems. These differences may reflect division rate, mutation-accumulation time, tissue environment or exogenous exposures, although the heterogeneous tumour cohort prevents attribution to any single factor. Consistent with this complexity, recent analyses of CMMRD tumours showed that mutational-signature composition depends on the defective MMR gene, tumour type, polymerase mutations and treatment history [35].

Finally, the noncanonical knockouts revealed additional repair phenotypes. EXO1 loss was enriched for SBS3/SBS8/SBS39 and DBS13, consistent with altered double-strand-break-repair pathway usage and with the established role of EXO1 in DNA-end resection and homologous recombination [23, 24]. Previous HAP1 experiments likewise identified substitution, indel and rearrangement phenotypes following EXO1 loss [18]. SETD2 loss was most clearly associated with SBS36, consistent with previous evidence linking SETD2-dependent H3K36me3 to DNA repair [36] and SETD2/MutS*α* to oxidative-damage processing [28]. SBS36 itself has been linked experimentally to defective base excision repair of 8-oxoG-associated lesions [19]. Although the study is limited by the single cell-line background, modest sequencing depth, small clone numbers and uncertainties inherent to NNLS refitting [37], the combined results broaden the mutational footprint of MMR deficiency and highlight the strong dependence of mutational signatures on biological and technical context.

## Methods

### Experimental strategy

We used CRISPR–Cas9 to create frameshift mutations in *MLH1*, *MSH2*, *MSH6*, *ARID1A*, *EXO1*, *SETD2*, *NSD1* and *SMARCA4* in the human HAP1 cell line. For each knockout cell line and a wildtype (WT) control, a single cell was selected from the early expansion pool and expanded for sequencing. These early-expansion cells were designated time point 0 (*t*_0_). The remaining cell pool was cultured for three months, after which three independently isolated single cells were expanded and sequenced as time point 1 (*t*_1_) clonal replicates. Most cells had become diploid by the end of the experiment. De novo variants were identified by paired somatic analysis of each *t*_1_ clone against its corresponding *t*_0_ sample.

For pooled analyses, the MMR-proficient (MMRp) group comprised ARID1A-sc1 and -sc2, NSD1sc1 and -sc2, SMARCA4-sc1, -sc2 and -sc3, and WT-sc1 (n = 8). The MMR-deficient (MMRd) group comprised MLH1-sc1, -sc4 and -sc5, MSH2-sc1, -sc2 and -sc3, and MSH6-sc1, -sc2 and -sc3 (n = 9). Per-clone mutation-burden and signature panels used controls WT, and three replicates each of ARID1A, EXO1, NSD1, SETD2, SMARCA4, MLH1, MSH2 and MSH6 samples.

### Cell culture

HAP1 cells were grown in Iscove’s Modified Dulbecco’s Medium (IMDM; GIBCO) containing Lglutamine and 25 mM HEPES and supplemented with 10% fetal bovine serum and 1% penicillin/streptomycin. Cells were grown at 37 *^○^*C with 5% oxygen and 5% carbon dioxide, passaged every 5 days, and kept subconfluent for 3 months. Cell lines tested negative for mycoplasma contamination using the MycoAlert mycoplasma detection kit. HAP1 is not listed in the ICLAC database of commonly misidentified cell lines. The parental HAP1 cell line was characterised and authenticated by Horizon Genomics. In our hands, HAP1 cells had a doubling time of approximately 24–48 h.

### Gene editing

CRISPR-Cas9 knockouts were generated in collaboration with Horizon Genomics. HAP1 cells were transfected with a Cas9-expression plasmid, a guide-RNA-expression plasmid and a plasmid conferring blasticidin resistance using Xfect (Clontech). Cells were treated with 20 *µ*g ml-1 blasticidin for 24 h to remove untransfected cells. After 5-7 days of recovery, clonal cell lines were isolated by limiting dilution. Guide-RNA sequences and cell lines are available from Horizon Genomics.

### Whole-genome sequencing (WGS) library preparation and sequencing

Short-insert paired-end whole-genome libraries were prepared using a PCR-free KAPA HyperPrep protocol (Roche), with modifications. One microgram of genomic DNA was sheared on a Covaris LE220-Plus and size-selected with AMPure XP beads (Beckman Coulter). DNA fragments were end-repaired and adenylated, and Illumina-compatible adaptors carrying unique dual indices and unique molecular identifiers (Integrated DNA Technologies) were ligated. Libraries were assessed on an Agilent 2100 Bioanalyzer using the DNA 7500 assay and quantified using the KAPA Library

#### Quantification Kit for Illumina platforms

Libraries were sequenced on an Illumina NovaSeq 6000 in paired-end mode (2 × 151 + 17 + 8 bp) following the manufacturer’s dual-indexing protocol. Image analysis, base calling and quality scoring used Real Time Analysis software version 3.4.4, followed by FASTQ generation. Sequencing was performed to a mean depth of approximately 15× per haploid genome. Ploidy was assessed by fluorescence-activated cell sorting before sequencing.

### Somatic variant calling

Cell-line reads were aligned to GRCh38 and analysed with nf-core/sarek version 3.8.1 in paired mode, with *t*_0_ specified as the normal sample and each *t*_1_ clone as the somatic sample [38]. Small variants were called with the Manta–Strelka2 and Mutect2 workflows. The relaxed “before-filtering” callset comprised de novo Strelka2 PASS variants in uniquely mappable sequence. Unique-mappability intervals on GRCh38 were derived from the 35-bp GEM mappability track hg38.fa.mappability_35bp.bw obtained from Peneder’s Zenodo mappability dataset (version 1.0.0; DOI: 10.5281/zenodo.5521424).

Only genomic positions contained within intervals having a 35-bp GEM mappability score of exactly 1.0 were considered uniquely mappable and included in the callable genome mask, while all other positions were excluded. The stringent cell-line “after-filtering” callset was obtained by additionally requiring concordant Strelka2 and Mutect2 PASS calls and an uppercase reference sequence at the variant-associated sequence to remove repeat-masked sequence.

### Mutational Signature Refitting

Mutational signatures were fitted by non-negative least squares using scipy.optimize.nnls, which implements the Lawson–Hanson algorithm, and the resulting NNLS signature activities reported. COSMIC version 3.4 references were used. SBS and DBS input profiles were derived directly from the data, while indel input profiles were generated with SigProfilerMatrixGenerator version 1.3.6 using the installed GRCh38 reference for cell lines and GRCh37 for tumours.

For Figure 2, the following single-base-substitution signatures were shown individually: SBS57, SBS7c, SBS1, SBS6, SBS14, SBS15, SBS20, SBS21, SBS26, SBS44, SBS17, SBS12 and SBS18. Other SBS signatures were grouped as follows: POLE, SBS10a/SBS10b/SBS28; POLD, SBS10c/SBS10d; DSB repair, SBS3/SBS8/SBS39; base-excision repair (BER), SBS30/SBS36; APOBEC, SBS2/SBS9/SBS13/ SBS84/SBS85; clock-like, SBS5/SBS40a/SBS40b/SBS40c; tobacco, SBS4/SBS29/SBS92; chemotherapy, SBS11/SBS25/SBS31/SBS35/SBS42/SBS86/SBS87/SBS99; environmental, SBS7a/SBS7b/SBS7d/ SBS22a/SBS22b/SBS24/SBS32/SBS88/SBS90; and all remaining signatures as unknown. For indel signatures, ID12, ID1, ID2 and ID7 were retained individually; ID6 and ID8 were grouped as DSB repair; ID17 was grouped as other repair; and all remaining ID signatures were grouped as unknown. For double-base-substitution signatures, DBS14, DBS16, DBS7 and DBS10 were retained individually; DBS3 was grouped as POLE, DBS13 as DSB repair, DBS2 as tobacco, DBS5 as chemotherapy, DBS1 and DBS20 as environmental, and all remaining DBS signatures were grouped as unknown. Display legends were restricted to categories with relevant activity in the corresponding panel. In other figures, SBS3/SBS30/SBS36 were grouped as Other repair, with SBS8 and SBS39 included in Unknown, ID6/ID8/ID17 were grouped as Other repair, and DBS13 was grouped as Other repair. SBS17a and SBS17b were combined as SBS17 in grouped panels but retained separately for the replication-timing slope analysis.

For the replication-time-dependent signature analyses, for each replication-timing decile *d*, the observed count *y_c,d_* in SBS96 channel *c* was divided by the callable opportunity *A_c,d_* for that channel and decile and multiplied by the corresponding fixed whole-genome opportunity *G_c_*. The resulting preliminary corrected total for decile d was calculated as *M_d_* = Σ*_c_ y_c,d_G_c_/A_c,d_*. A single scaling factor, calculated as the total observed mutation count across all deciles divided by Σ*_d_ M_d_*, was applied to every channel and decile, preserving the overall mutation total while retaining opportunity-corrected differences among deciles. Each corrected decile-specific profile was then fitted independently by NNLS. The plotted values are the resulting NNLS activities divided by the number of samples in the corresponding group.

The TensorSignatures analysis used published source-data spectra from Vöhringer et al. [12]. The 33-column reference comprised TS01, TS02, TS03.1, TS03.2, TS04, TS05–TS09, TS10-a, TS10-q, TS11, TS12a, TS12b and TS13–TS30. For each source signature, an unstranded SBS96 profile was derived from the coding, template, leading and lagging marginal spectra. The coding-plus-template vector and leading-plus-lagging vector were each normalised to sum to one, their arithmetic mean was then calculated and renormalised to unit sum.

The same RT bin assignment, callable-opportunity calculation, whole-genome scaling and global count-preserving rescaling used for the COSMIC analysis were applied to the TensorSignatures analysis. NNLS was then performed separately for MMRp-before, MMRd-before, MMRp-after and MMRd-after profiles, and activities were divided by the corresponding number of samples.

### Tumour data

Tumour analyses used HMF metastatic-tumour calls in GRCh37/hg19 coordinates [32], read from PURPLE somatic VCF files. Variant records were retained when FILTER was PASS or “.”, the chromosome was 1–22, X or Y, and the variant was a biallelic A/C/G/T SNV or a length-changing indel. For the tumour “before-filtering” tables, variants were restricted to uniquely mappable sequence. Uniquely mappable intervals were derived from the CRG 36-mer alignability track wgEncodeCrgMapabilityAlign36mer.bigWig obtained from the UCSC ENCODE archive. Only intervals with a mappability score of exactly 1.0 were retained. The tumour “after-filtering” tables additionally required uppercase reference sequence at the variant-associated sequence, thereby excluding repeat-masked sequence.

For the final after-filtering SNV callsets, variants near deletions were removed in a sample- and chromosome-specific manner. For each deletion, the deleted interval from event_start through event-_start + deleted length 1 was expanded by 6 bp on both sides. Overlapping windows were merged, and an SNV was removed when its 1-based genomic position fell within a merged window. This procedure was applied independently to the MMRd-like, SBS17-like and Other tumour groups, removing 1.4% of variants for the MMRd tumour cluster, 0.15% for the SBS17 cluster and 0.3% for the Other cluster. For **Supplementary Figure 6**, an alternative MMRd tumour after-filtering input retaining near-deletion SNVs was analysed.

### Replication timing

To determine replication time, we computed the mean wavelet-smoothed signal across 6 cell lines measured by the ENCODE project [39]. The cell lines are Gm12878, Helas3, Hepg2, Huvec, K562, and Mcf7. The data files and download locations are documented at https://genome.ucsc.edu/cgibin/hgTrackUi?db=hg19&g=wgEncodeUwRepliSeq. Replication-timing values were originally defined on hg19, and the corresponding intervals were lifted to hg38 and restricted to primary chromosomes for analysis of the cell-line data. Higher scores represent earlier replication. Sites without RT coverage were excluded from RT-resolved analyses.

Callable trinucleotide opportunities were counted directly from chromosome FASTA sequences for each RT bin. The before-filtering opportunity mask required RT coverage, mappability and a valid A/C/G/T trinucleotide, accepting both upperand lowercase bases. The after-filtering mask additionally required an uppercase central base. Whole-genome trinucleotide abundances were computed directly from the reference FASTA sequences, without applying replication-timing, mappability, or lowercase-sequence filters.

**Figure 4D** used the after-filtering RT-resolved activities for SBS1, SBS6, SBS14, SBS15, SBS20, SBS21, SBS26, SBS44, SBS17a, SBS17b and SBS18. Deciles were encoded from 1 (earliest) to 10 (latest), and a first-degree polynomial was fitted with numpy.polyfit for each signature and group. The reported slope therefore has units of average assigned mutations per sample per RT decile, with positive values indicating increasing activity towards late replication.

### Replication and transcription strand-bias analyses

Strand labels were recomputed from genomic positions and external annotations. SBS mutations were oriented to the pyrimidine-mutated strand. Cell-line analyses used the GRCh38 six-cell-line RT track and GENCODE v49 GRCh38 gene annotations, while tumour analyses used the hg19 RT track and GENCODE v49lift37 annotations.

Replication-fork direction was inferred from the local RT-score gradient. For internal intervals, the gradient was score(i+1) score(i-1) and endpoint intervals used the adjacent score difference. Because higher scores indicate earlier replication, a negative gradient was assigned a fork moving in the positive genomic direction and a positive gradient a fork moving in the negative direction. Gradients with absolute value ≤ 10^—12^ were considered ambiguous. For a positive-direction fork, a mutation represented on the genomic minus pyrimidine strand was labelled leading and one on the plus strand lagging. The assignments were reversed for a negative-direction fork. Uncovered and ambiguous sites were excluded from strand-specific counts.

For transcription strand, only GENCODE gene features on chromosomes 1–22, X and Y were used. A mutation overlapping only plus-strand genes was labelled transcribed when its pyrimidine-mutated strand was genomic minus, while a mutation overlapping only minus-strand genes was labelled transcribed when its pyrimidine-mutated strand was genomic plus. Intergenic mutations and mutations overlapping genes on both strands were unassigned and excluded from transcription-bias counts. Strand-specific SBS96 counts were summed within each group and divided by the number of samples.

### Statistical analysis

Enrichment of individual or grouped signature activities in EXO1 or SETD2 knockout clones relative to the MMRp group was assessed using a two-sided exact permutation test based on the difference in group means. For each comparison, the activities from the three target-gene knockout clones and eight MMRp samples were pooled, and all 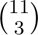 =165 possible assignments of three samples to the target group were enumerated. The exact P-value was the proportion of permutations for which the absolute difference in means was at least as large as the observed value. Fold enrichment was calculated as the target-group mean divided by the MMRp-group mean. Reported P-values are raw P-values, not corrected for multiple testing.

### SBS57-positive tumour selection, principal-component analysis and clustering

To estimate the mutation-burden-dependent rate of false SBS57 assignment by non-negative leastsquares fitting, we constructed an SBS57-negative simulated null distribution from 2,780 PCAWG signature-activity profiles having no assigned SBS57 contribution [40]. PCAWG source spectra were reconstructed using the COSMIC version 3.3.1 GRCh37 SBS reference. For each source profile, SBS96 catalogues containing 25, 50, 100, 250, 500, 1,000, 5,000, 10,000, 50,000 or 100,000 mutations were generated by multinomial sampling, with ten replicate catalogues per source profile and mutation burden. This produced 278,000 simulated catalogues with no true SBS57 contribution. Each simulated catalogue was refitted by non-negative least squares using the COSMIC version 3.4 GRCh37 SBS reference, including SBS57. For each simulated mutation burden, the upper 0.1% quantile of the fitted SBS57 relative weight was calculated. The burden-specific thresholds were constrained to be monotonically non-increasing with mutation burden using isotonic regression and were interpolated on the logarithm of mutation burden.

COSMIC version 3.4 SBS signatures were subsequently fitted to 4,709 HMF SBS96 catalogues. HMF tumours were selected when they contained at least 1,000 SBS mutations and their fitted SBS57 relative weight exceeded the interpolated, burden-dependent 99.9th-percentile threshold derived from the SBS57-negative simulated null distribution. This procedure identified 223 SBS57-positive HMF tumours.

Principal-component analysis was performed on centred, but not variance-scaled, COSMIC signature-weight profiles. SBS57 was excluded from the PCA features so that clustering was not driven directly by the signature used for tumour selection. The first two principal components were used for k-means clustering. Candidate solutions with k=2 through k=8 were compared by average silhouette width and k=3 gave the highest average silhouette width (0.63).

The cluster with the largest mean combined contribution from SBS6, SBS14, SBS15, SBS20, SBS21, SBS26 and SBS44 was labelled MMRd-like. Of the remaining clusters, the cluster with the largest mean combined SBS17a and SBS17b contribution was labelled SBS17-like, and the remaining cluster was labelled Other. The three clusters initially contained 42 MMRd-like, 87 SBS17-like and 94 Other tumours.

Samples were considered robustly assigned when their individual silhouette width exceeded the mean silhouette width of the three-cluster solution and their cluster contained at least five such samples. This retained 29 MMRd-like, 48 SBS17-like and 68 Other tumour samples. Clustering and silhouette calculations were performed at the tumour-sample level. For downstream analyses, one sample per patient was retained, preferring an exact primary T sample and otherwise the earliest retained longitudinal sample. This produced patient-level groups of 29 MMRd-like, 47 SBS17-like and 66 Other tumours.

## Data availability

Sequencing data generated for this project will be made publicly available. Pan-cancer mutation signature activities were retrieved from the PCAWG (Pan-Cancer Analysis of Whole Genomes) database (docs.icgc-argo.org/docs/data-access/icgc-25k-data). Restricted-access data from the Hartwig Medical Foundation were obtained after application from www.hartwigmedicalfoundation.nl/en/.

## Acknowledgments

We acknowledge support of the Spanish Ministry of Science and Innovation through the Centro de Excelencia Severo Ochoa (CEX2020-001049-S, MCIN/AEI /10.13039/501100011033), and the Generalitat de Catalunya through the CERCA programme. This work was further supported by grant PID2021-128976NB-I00 to D.W. funded by MICIU/AEI/10.13039/501100011033/FEDER, UE, and by the FWF Austrian Science Fund with an Erwin-Schrödinger postdoctoral fellowship, No. J4366 (to M.O.). This publication and the underlying study have been made possible partly based on data that Hartwig Medical Foundation and the Center of Personalised Cancer Treatment (CPCT) have made available to the study through the Hartwig Medical Database.

## Author contributions

Conceptualisation: M.O., D.W., data curation: M.O., D.W., formal analysis: M.O., D.W., investigation: M.O., D.W., methodology: M.O., J.L., D.W., resources: J.M., J.L., D.W., supervision: D.W., writing: M.O., D.W., funding acquisition: M.O., J.M., D.W.

## References

[1] Su, S.-S. & Modrich, P. Escherichia coli muts-encoded protein binds to mismatched dna base pairs. Proceedings of the National Academy of Sciences 83, 5057–5061 (1986).

[2] Liu, D., Keijzers, G. & Rasmussen, L. J. Dna mismatch repair and its many roles in eukaryotic cells. Mutation Research/Reviews in Mutation Research 773, 174–187 (2017).

[3] Meier, B. et al. Mutational signatures of DNA mismatch repair deficiency in C. elegans and human cancers. Genome research 28, 666–675 (2018).

[4] Fang, H. et al. Deficiency of replication-independent dna mismatch repair drives a 5methylcytosine deamination mutational signature in cancer. Science Advances 7, eabg4398 (2021).

[5] Poulos, R. C., Olivier, J. & Wong, J. W. The interaction between cytosine methylation and processes of dna replication and repair shape the mutational landscape of cancer genomes. Nucleic acids research 45, 7786–7795 (2017).

[6] Aaltonen, L. A. et al. Clues to the pathogenesis of familial colorectal cancer. Science 260, 812–816 (1993).

[7] Wang, Q. et al. Neurofibromatosis and early onset of cancers in h mlh1-deficient children. Cancer Research 59, 294–297 (1999).

[8] Shlien, A. et al. Combined hereditary and somatic mutations of replication error repair genes result in rapid onset of ultra-hypermutated cancers. Nature genetics 47, 257–262 (2015).

[9] Andrianova, M. A. et al. Germline pms2 and somatic pole exonuclease mutations cause hypermutability of the leading dna strand in biallelic mismatch repair deficiency syndrome brain tumours. The Journal of pathology 243, 331–341 (2017).

[10] Baretti, M. & Le, D. T. Dna mismatch repair in cancer. Pharmacology & therapeutics 189, 45–62 (2018).

[11] Alexandrov, L. B. et al. The repertoire of mutational signatures in human cancer. Nature 578, 94–101 (2020).

[12] Vöhringer, H., Van Hoeck, A., Cuppen, E. & Gerstung, M. Learning mutational signatures and their multidimensional genomic properties with tensorsignatures. Nature Communications 12, 1–16 (2021).

[13] Lowenthal, B. M. et al. Loss of ARID1A expression is associated with DNA mismatch repair protein deficiency and favorable prognosis in advanced stage surgically resected esophageal adenocarcinoma. Human Pathology 94, 1–10 (2019).

[14] Genschel, J. & Modrich, P. Mechanism of 5’-Directed Excision in Human Mismatch Repair. Molecular Cell 12, 1077–1086 (2003).

[15] Li, F. et al. The Histone Mark H3K36me3 Regulates Human DNA Mismatch Repair through Its Interaction with MutS*α*. Cell 153, 590–600 (2013).

[16] Li, F., Ortega, J., Gu, L. & Li, G.-M. Regulation of mismatch repair by histone code and posttranslational modifications in eukaryotic cells. DNA Repair 38, 68–74 (2016).

[17] Drost, J. et al. Use of crispr-modified human stem cell organoids to study the origin of mutational signatures in cancer. Science 358, 234–238 (2017).

[18] Zou, X. et al. Validating the concept of mutational signatures with isogenic cell models. Nature communications 9, 1–16 (2018).

[19] Zou, X. et al. A systematic crispr screen defines mutational mechanisms underpinning signatures caused by replication errors and endogenous dna damage. Nature cancer 2, 643–657 (2021).

[20] Colussi, C. et al. The mammalian mismatch repair pathway removes dna 8-oxodgmp incorporated from the oxidized dntp pool. Current biology 12, 912–918 (2002).

[21] Martin, S. A. et al. Methotrexate induces oxidative dna damage and is selectively lethal to tumour cells with defects in the dna mismatch repair gene msh2. EMBO molecular medicine 1, 323–337 (2009).

[22] Ohno, M. et al. Oxidative stress accelerates repeat sequence instability and base substitutions promoting gastrointestinal driver mutations in msh2 deficient mice. Genes and Environment 47, 19 (2025).

[23] Tomimatsu, N. et al. Exo1 plays a major role in dna end resection in humans and influences double-strand break repair and damage signaling decisions. DNA repair 11, 441–448 (2012).

[24] Bolderson, E. et al. Phosphorylation of exo1 modulates homologous recombination repair of dna double-strand breaks. Nucleic acids research 38, 1821–1831 (2010).

[25] Singh, V. K., Rastogi, A., Hu, X., Wang, Y. & De, S. Mutational signature sbs8 predominantly arises due to late replication errors in cancer. Communications biology 3, 421 (2020).

[26] Degasperi, A. et al. A practical framework and online tool for mutational signature analyses show intertissue variation and driver dependencies. Nature cancer 1, 249–263 (2020).

[27] Degasperi, A. et al. Substitution mutational signatures in whole-genome–sequenced cancers in the UK population. Science 376 (2022).

[28] Guo, S. et al. Interplay between h3k36me3, methyltransferase setd2, and mismatch recognition protein muts*α* facilitates processing of oxidative dna damage in human cells. Journal of Biological Chemistry 298 (2022).

[29] Basyuni, S. et al. Large-scale analysis of whole genome sequencing data from formalin-fixed paraffin-embedded cancer specimens demonstrates preservation of clinical utility. Nature Communications 15, 7731 (2024).

[30] Delhomme, T. M. et al. Proton and alpha radiation-induced mutational profiles in human cells. Scientific Reports 13, 9791 (2023).

[31] Everall, A. et al. Comprehensive repertoire of the chromosomal alteration and mutational signatures across 16 cancer types. Nature Genetics 58, 570–581 (2026).

[32] Priestley, P. et al. Pan-cancer whole-genome analyses of metastatic solid tumours. Nature 575, 210–216 (2019).

[33] Spisak, N., de Manuel, M., Milligan, W., Sella, G. & Przeworski, M. The clock-like accumulation of germline and somatic mutations can arise from the interplay of dna damage and repair. PLoS biology 22, e3002678 (2024).

[34] Cosmic mutational signatures (v3.4 october 2023). gnatures/sbs/. Accessed: 2023-10-24.

[35] Weijers, D. D. et al. Unraveling mutagenic processes influencing the tumor mutational patterns of individuals with constitutional mismatch repair deficiency. Nature Communications 16, 4459 (2025).

[36] Pfister, S. X. et al. Setd2-dependent histone h3k36 trimethylation is required for homologous recombination repair and genome stability. Cell reports 7, 2006–2018 (2014).

[37] Serrano Colome, C., Canal Anton, O., Seplyarskiy, V. & Weghorn, D. Mutational signature decomposition with deep neural networks reveals origins of clock-like processes and hypoxia dependencies. bioRxiv (2023). Publisher: Cold Spring Harbor Laboratory _eprint: https://www.biorxiv.org/content/early/2023/12/08/2023.12.06.570467.full.pdf.

[38] Hanssen, F., et al. Scalable and efficient dna sequencing analysis on different compute infrastructures aiding variant discovery. NAR Genomics and Bioinformatics 6, lqae031 (2024).

[39] Davis, C. A. et al. The encyclopedia of dna elements (encode): data portal update. Nucleic acids research 46, D794–D801 (2018).

[40] The ICGC/TCGA Pan-Cancer Analysis of Whole Genomes Consortium. Pan-cancer analysis of whole genomes. Nature 578, 82–93 (2020).

