## Supplementary Figures for "Dissecting DNA-mismatch-repair-driven mutational processes in human cells"

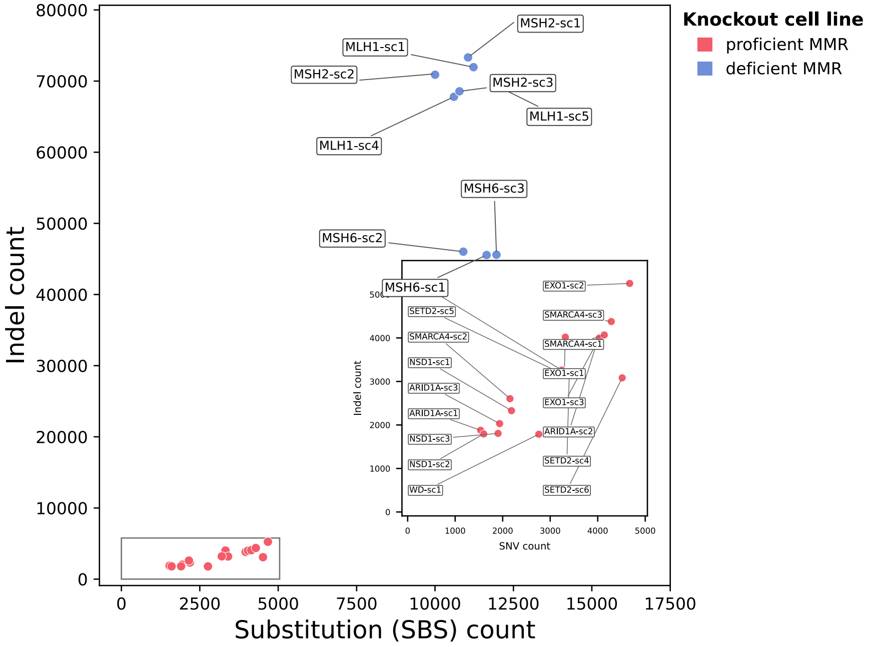


**Supplementary Figure 1:** Counts of *de novo* single-base substitutions and small insertions and deletions (indels) are shown for all KOs (including technical replicates) and wild-type cells (WT) for the "before-filtering" case, i.e., after Strelka2-based variant calling and restriction to uniquely mappable sites.

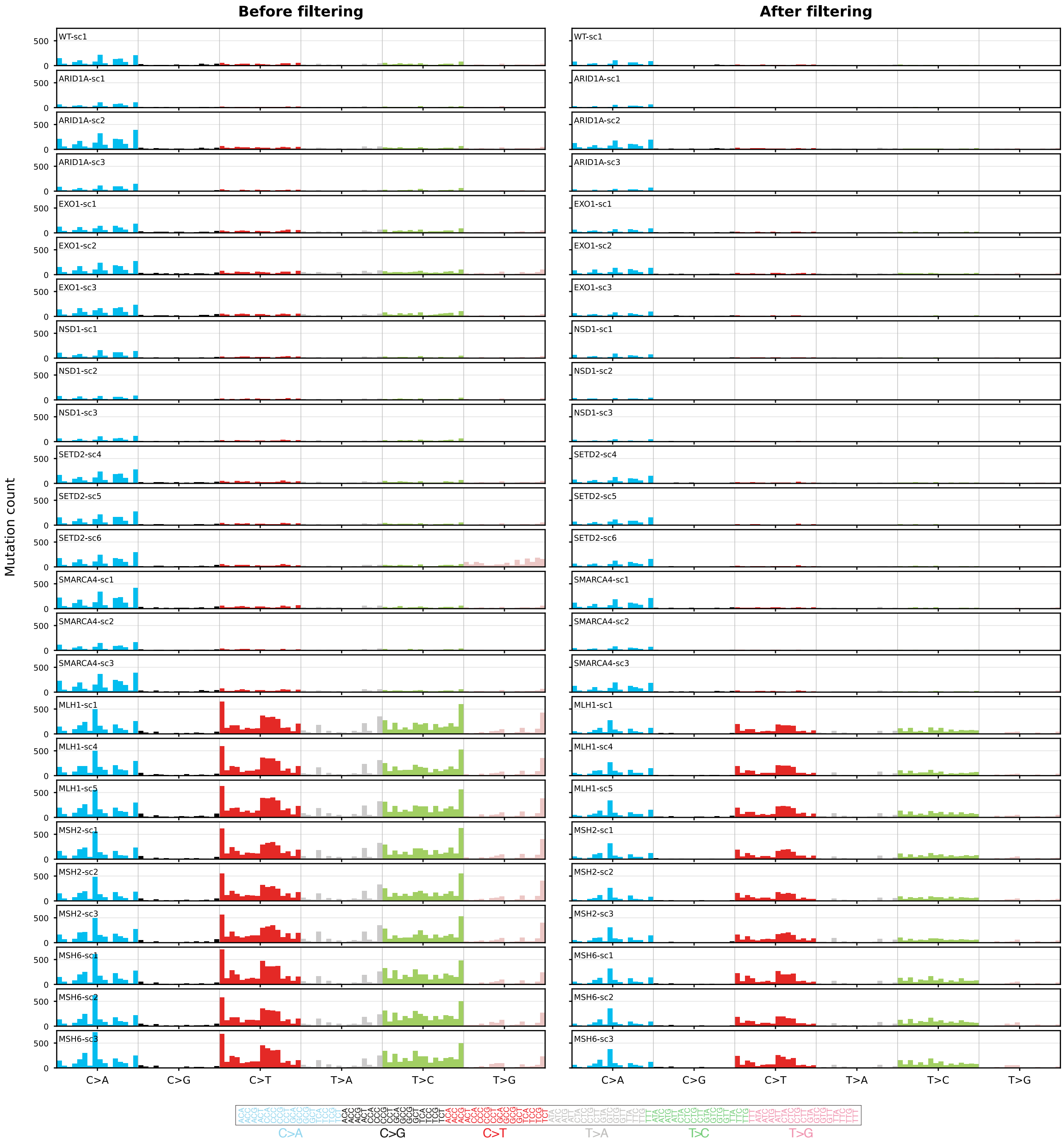


**Supplementary Figure 2:** Single-base-substitution (SBS) mutational profiles of all cell line knockouts, including technical replicates, before filtering (left) and after filtering (right). Before-filtering data were obtained from Strelka2 variant calling and restriction to uniquely mappable sites. After-filtering data were obtained by additionally requiring intersection with Mutect2 variant calling and restriction to non-repeat-masked genomic regions.

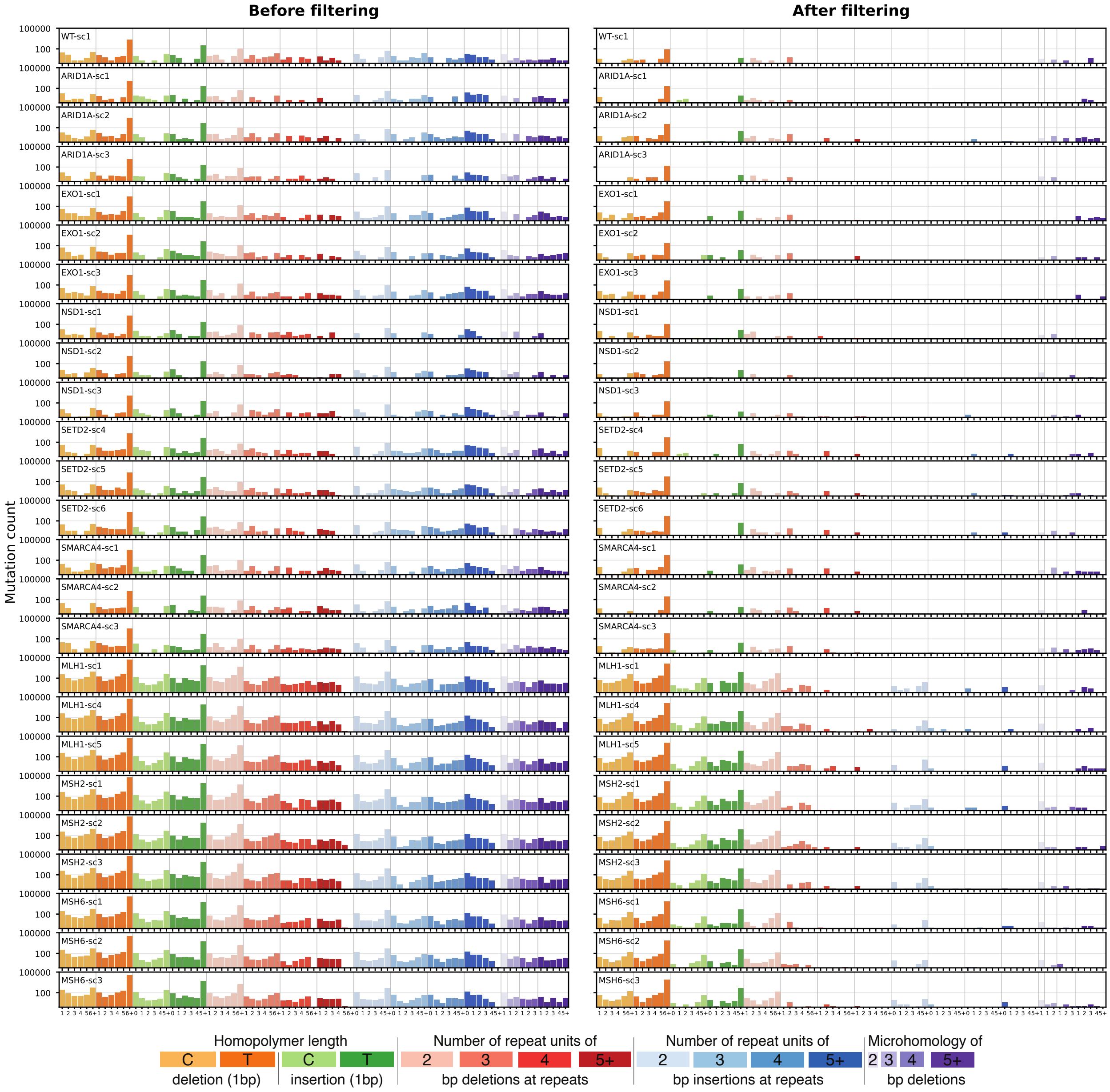


**Supplementary Figure 3:** Small insertion and deletion (ID) mutational profiles of all cell line knockouts, including technical replicates, before filtering (left) and after filtering (right). Before-filtering data were obtained from Strelka2 variant calling and restriction to uniquely mappable sites. After-filtering data were obtained by additionally requiring intersection with Mutect2 variant calling and restriction to non-repeat-masked genomic regions. Note the logarithmic y-axis.

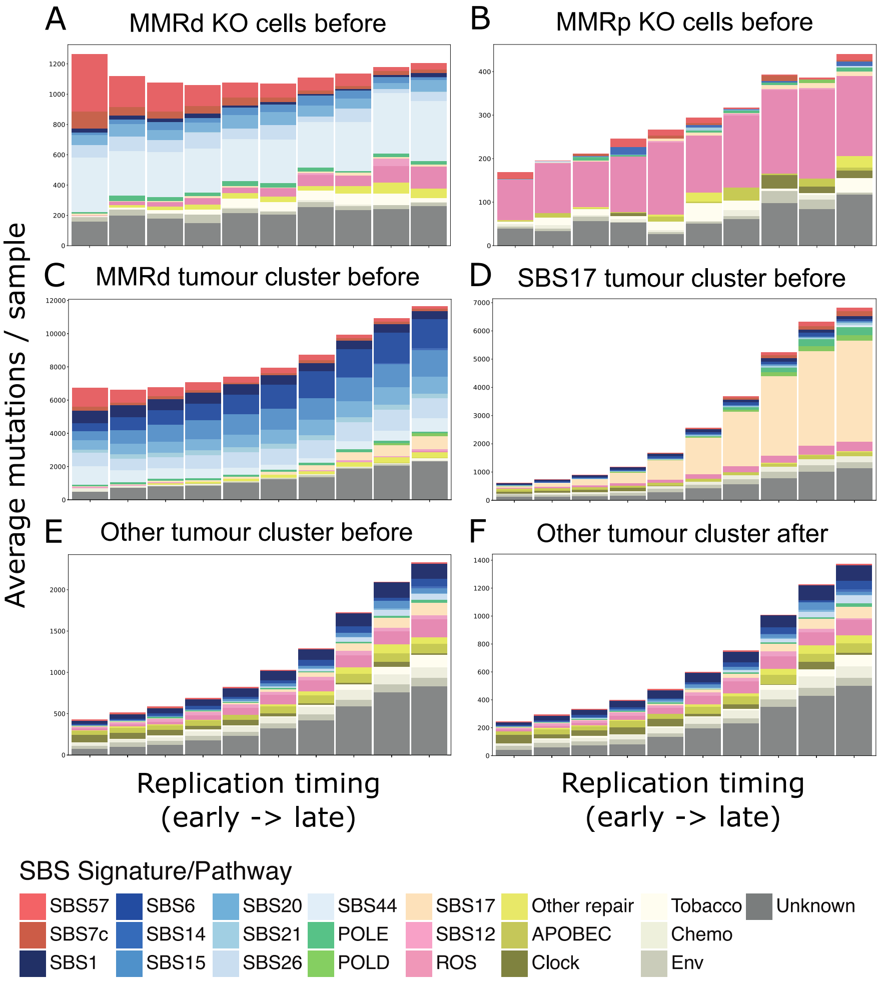


**Supplementary Figure 4:** Sequence-context-opportunity-corrected replication time dependence of SBS-signature-associated mutation counts in (A) MMRd cells before additional filtering, (B) MMRp cells before additional filtering, (C) MMRd-like tumours before additional filtering, (D) SBS17-like tumours before additional filtering, (E) Other tumours before additional filtering and (F) Other tumours after additional filtering, obtained from non-negative least-squares refitting to COSMIC v3.4 signatures. For cells, before-filtering data are Strelka2 variant calls restricted to uniquely mappable sites; after-filtering data additionally require intersection with Mutect2 variant calls and exclude variants in repeat-masked regions. For tumours, before-filtering data are HMF-provided Strelka calls restricted to uniquely mappable sites; after-filtering data exclude variants in repeat-masked regions and/or within ±6 bp of small deletions.

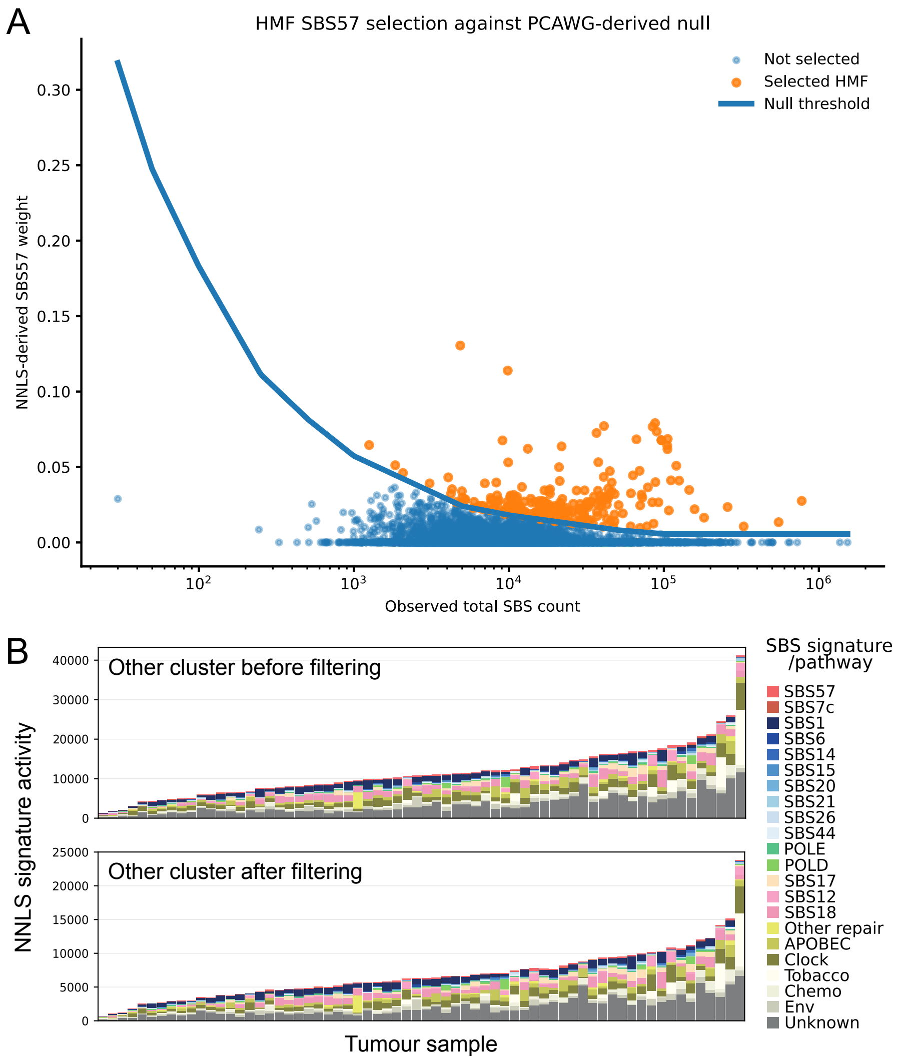


**Supplementary Figure 5:** (A) Selection of SBS57-positive tumours from 4709 HMF mutational profiles. Each point represents one tumour and shows its observed SBS burden and the relative SBS57 weight obtained by non-negative least-squares refitting to COSMIC v3.4 signatures. The solid line denotes the mutation-burden-dependent 99.9th-percentile threshold derived from simulated SBS57-negative mutational profiles based on PCAWG-predicted signature activities. HMF tumours with at least 1,000 SBSs and an SBS57 weight above this threshold were classified as SBS57-positive (orange; n=223); all other tumours are shown in blue. (B) Signature refitting of SBS57-containing tumour samples from the Other cluster. Top: Mutation calling without extra filters (i.e., HMF-provided Strelka calls restricted to mappable genomic regions). Bottom: Mutation calling with applying the additional filtering scheme (i.e., exclusion of variants in repeat-masked regions and/or within ±6 bp of small deletions). HMF=Hartwig Medical Foundation.

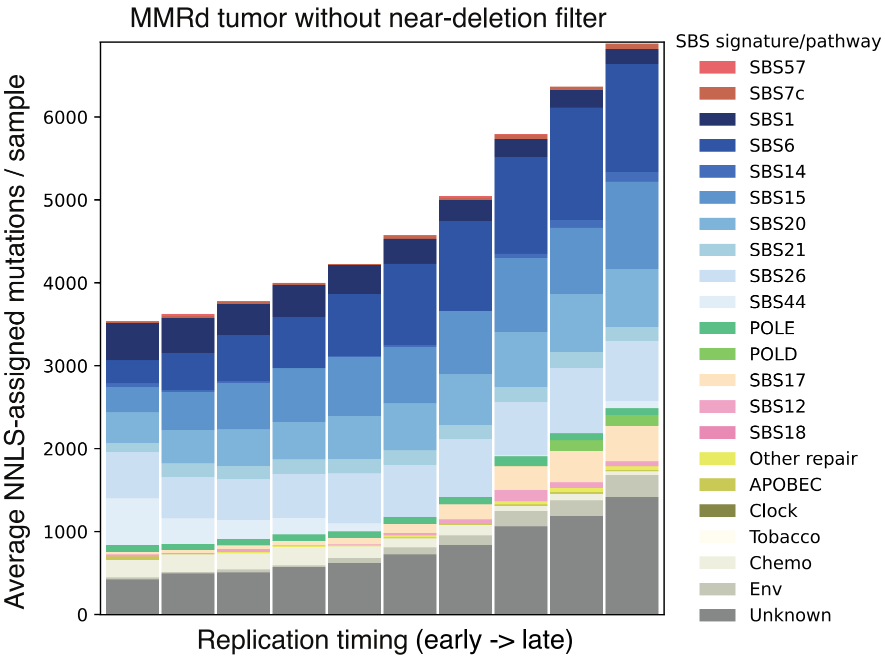


**Supplementary Figure 6:** Non-negative least-squares refitting of SBS57-containing tumour samples from the MMRd-like cluster to COSMIC v3.4 signatures, using HMF-provided Strelka calls restricted to uniquely mappable genomic regions and excluding variants in repeat-masked regions. Variants within ±6 bp of small deletions were included.

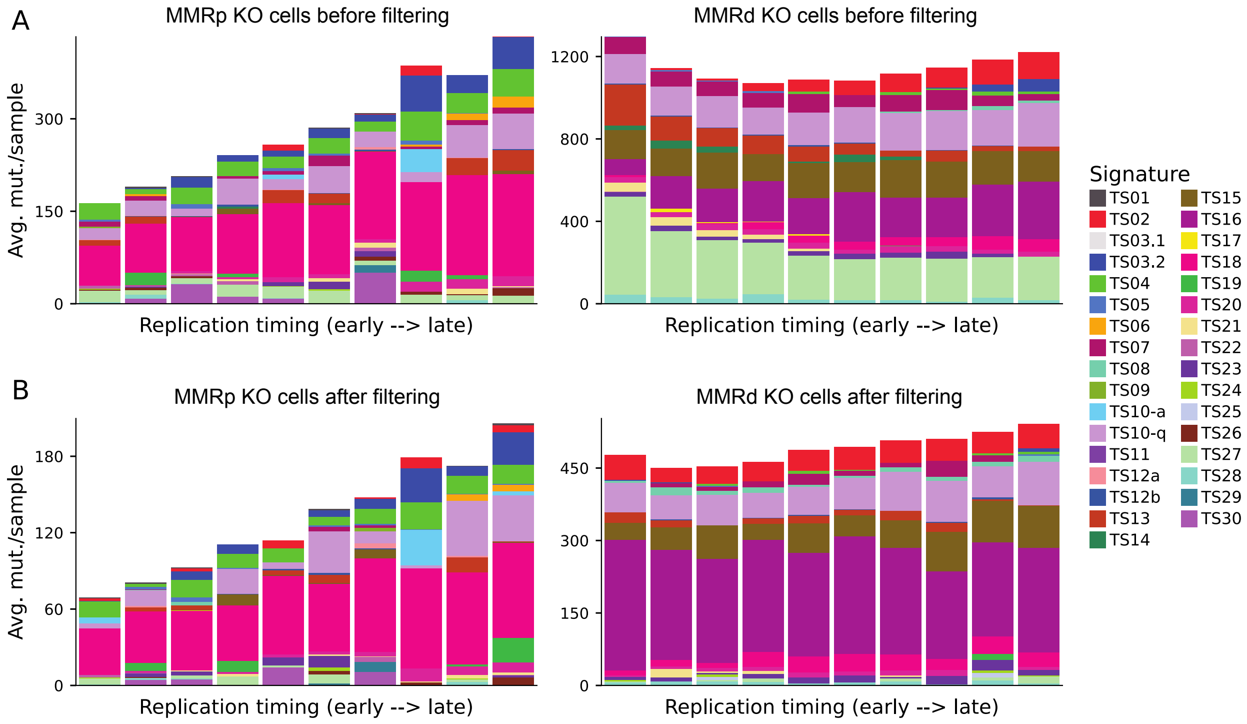


**Supplementary Figure 7:** Sequence-context-opportunity-corrected replication time dependence of TensorSignatures-associated mutation counts in MMRp (left) and MMRd HAP1 cells (right) obtained from non-negative least-squares refitting to tensor signatures [12]. (A) Before-filtering data: Strelka2 variant calls restricted to uniquely mappable sites. (B) After-filtering data: Additional intersection with Mutect2 variant calls and excluding variants in repeat-masked regions.

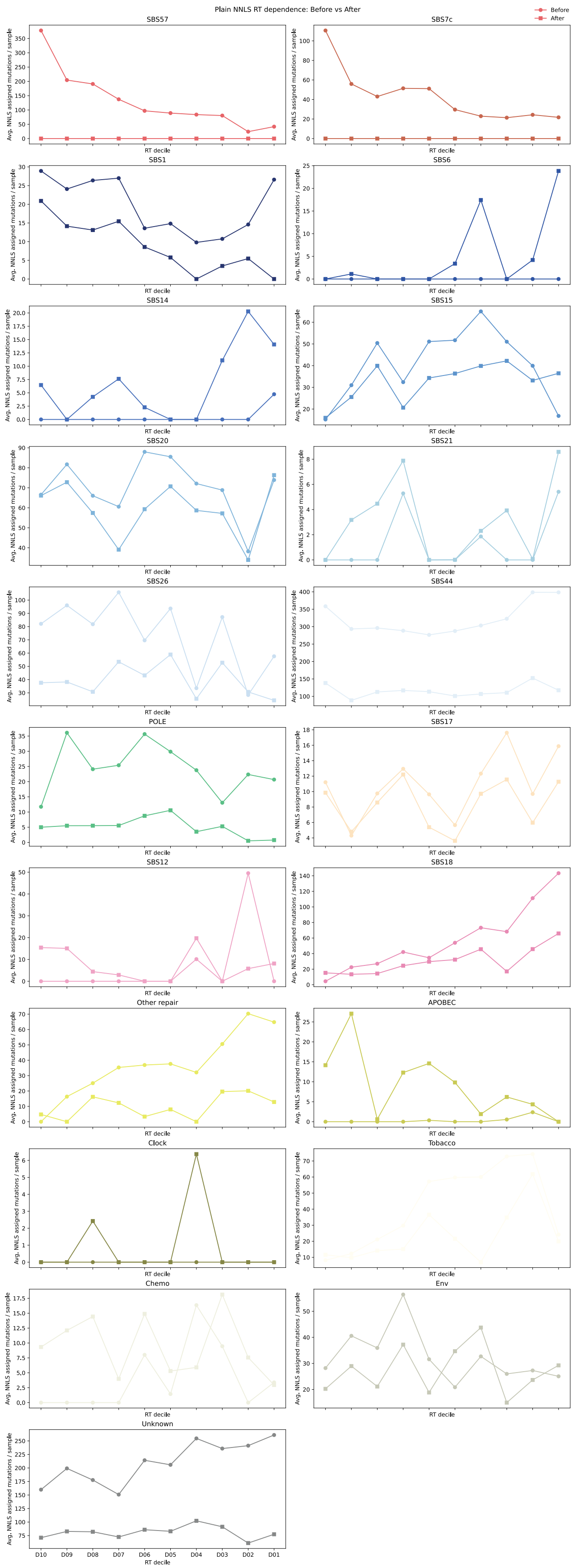


**Supplementary Figure 8:** Sequence-context-opportunity-corrected replication time dependence of COSMIC mutational signatures for MMRd cells before and after filtering.

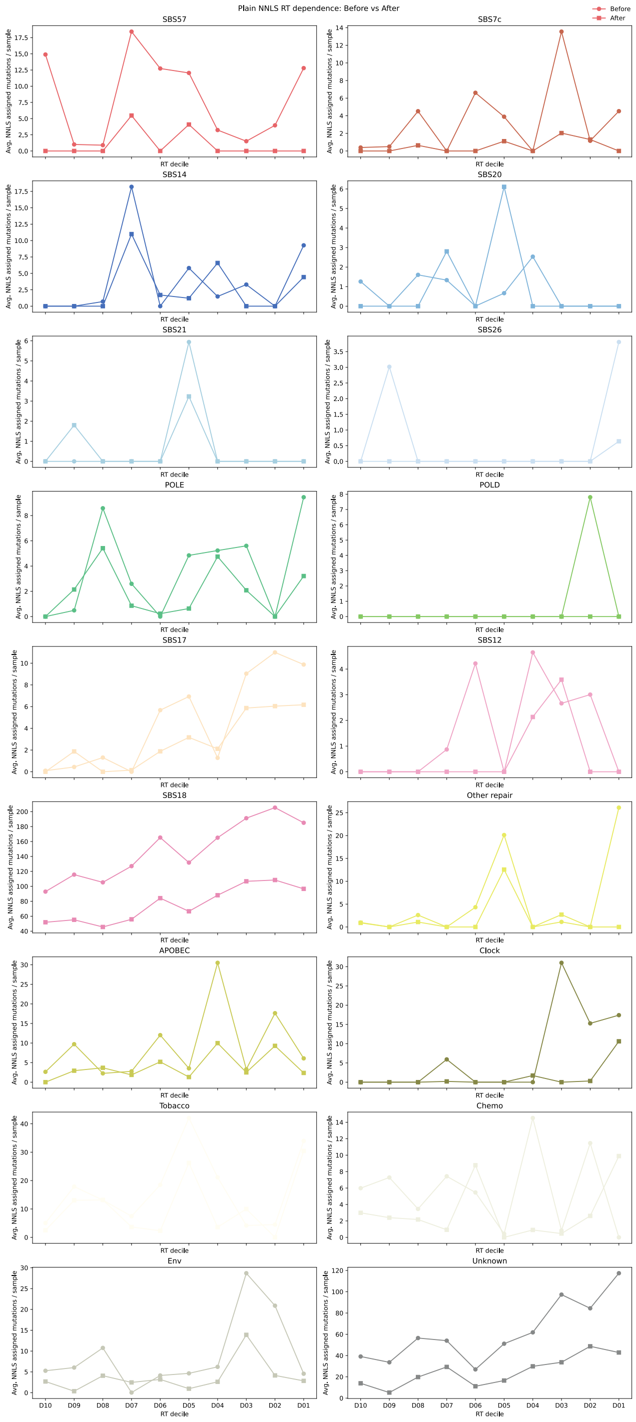


**Supplementary Figure 9:** Sequence-context-opportunity-corrected replication time dependence of COSMIC mutational signatures for MMRp cells before and after filtering.

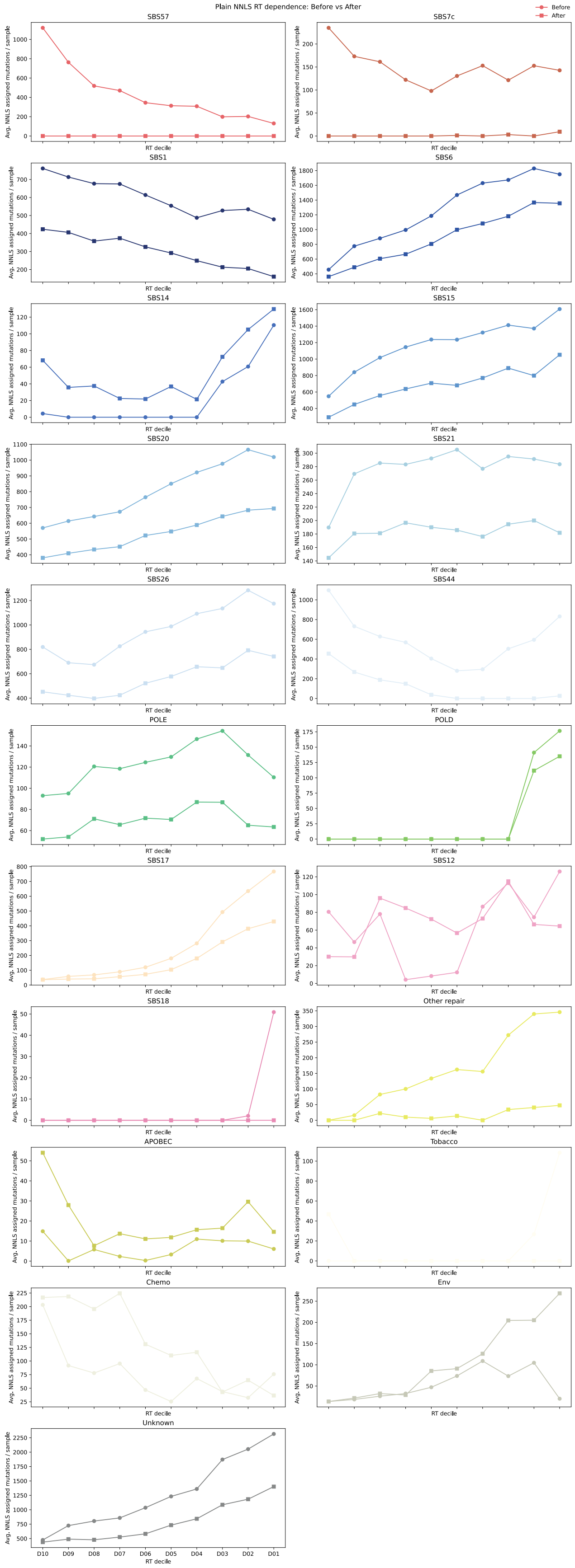


**Supplementary Figure 10:** Sequence-context-opportunity-corrected replication time dependence of COSMIC mutational signatures for MMRd-like tumours before and after filtering.

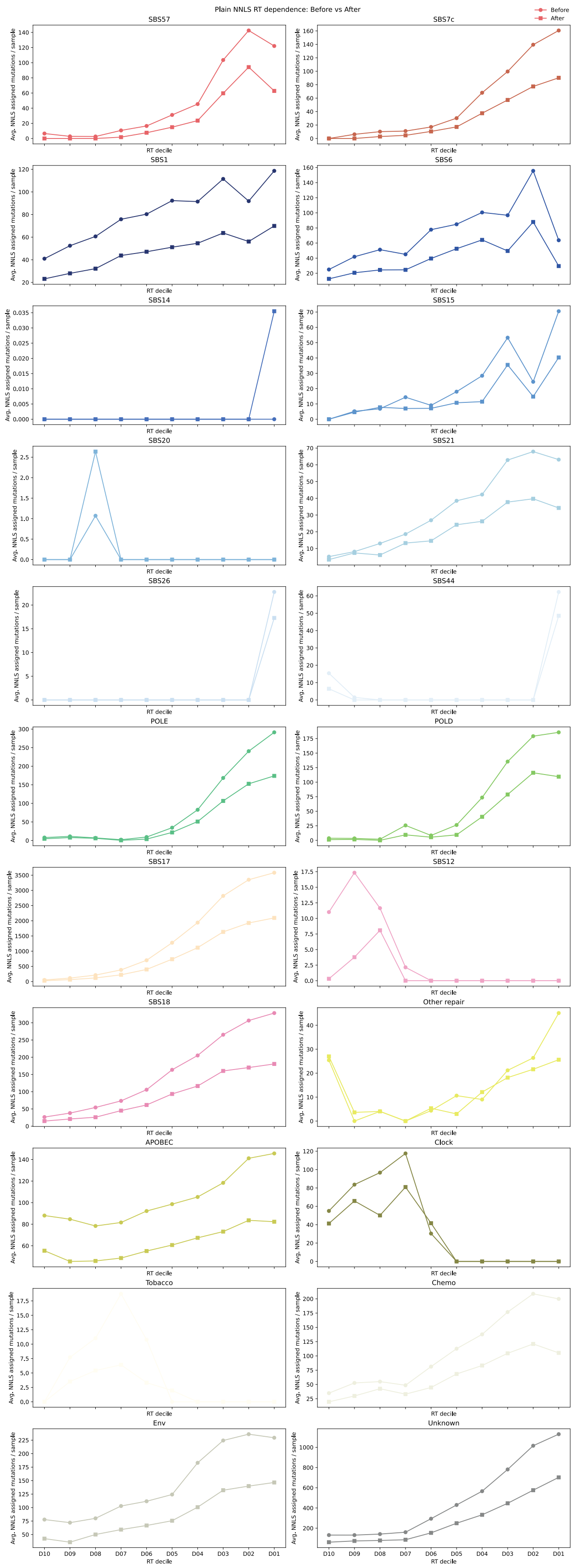


**Supplementary Figure 11:** Sequence-context-opportunity-corrected replication time dependence of COSMIC mutational signatures for SBS17-like tumours before and after filtering.

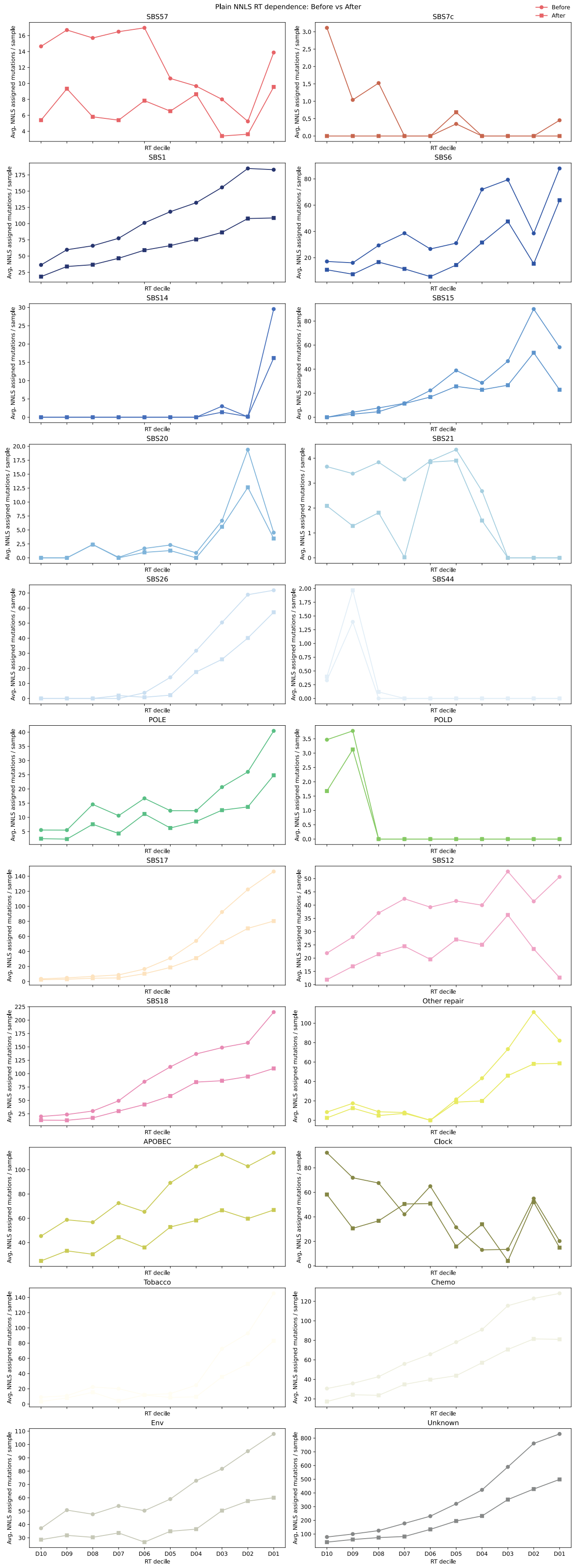


**Supplementary Figure 12:** Sequence-context-opportunity-corrected replication time dependence of COSMIC mutational signatures for Other SBS57-containing tumours before and after filtering.
